# What, if anything, is the true neurophysiological significance of “rotational dynamics”?

**DOI:** 10.1101/597419

**Authors:** Mikhail A. Lebedev, Alexei Ossadtchi, Nil Adell Mill, Núria Armengol Urpí, Maria R. Cervera, Miguel A.L. Nicolelis

**Affiliations:** Duke Center for Neuroengineering, Duke University, Durham, NC, USA; Center for Bioelectric Interfaces of the Institute for Cognitive Neuroscience of the National Research University Higher School of Economics, Moscow, Russia; Department of Information and Internet Technologies of Digital Health Institute, I.M. Sechenov First Moscow State Medical University, Moscow, Russia; Institute of Neuroinformatics, University of Zurich and ETH Zurich, Switzerland; D-MAVT ETH Zurich, Switzerland; Department of Neurobiology, Duke University Medical Center, Durham, NC, USA; Department of Neurology, Duke University, Durham, NC, USA; Department of Neurosurgery, Duke University, Durham, NC, USA; Department of Psychology and Neuroscience, Duke University, Durham, NC, USA; Edmond and Lily Safra International Institute of Neurosciences of Natal, Natal, Brazil

## Abstract

Back in 2012, Churchland and his colleagues proposed that “rotational dynamics”, uncovered through linear transformations of multidimensional neuronal data, represent a fundamental type of neuronal population processing in a variety of organisms, from the isolated leech central nervous system to the primate motor cortex. Here, we evaluated this claim using Churchland’s own data and simple simulations of neuronal responses. We observed that rotational patterns occurred in neuronal populations when (1) there was a temporal shift in peak firing rates exhibited by individual neurons, and (2) the temporal sequence of peak rates remained consistent across different experimental conditions. Provided that such a temporal order of peak firing rates existed, rotational patterns could be easily obtained using a rather arbitrary computer simulation of neural activity; modeling of any realistic properties of motor cortical responses was not needed. Additionally, arbitrary traces, such as Lissajous curves, could be easily obtained from Churchland’s data with multiple linear regression. While these observations suggest that temporal sequences of neuronal responses could be visualized as rotations with various methods, we express doubt about Churchland et al.’s exaggerated assessment that such rotations are related to “an unexpected yet surprisingly simple structure in the population response”, which “explains many of the confusing features of individual neural responses.” Instead, we argue that their approach provides little, if any, insight on the underlying neuronal mechanisms employed by neuronal ensembles to encode motor behaviors in any species.

## Introduction

It is well known that individual neurons in cortical motor areas transiently modulate their firing rates following a stimulus that triggers the production of a voluntary movement (Evarts 1972). These neuronal modulations have been shown to represent various motor parameters, for example movement direction (Georgopoulos, Kalaska et al. 1982), although the specifics of these representations are still a matter of debate (Georgopoulos, Ashe et al. 1992, Kakei, Hoffman et al. 1999, Zhuang, Lebedev et al. 2014). With the development of multichannel recordings (Nicolelis, Dimitrov et al. 2003, Schwarz, Lebedev et al. 2014), it has become possible to study modulations recorded in multiple cortical neurons simultaneously. This methodological advance led to many studies attempting to uncover how neuronal populations process information (Chapin and Nicolelis 1999, Laubach, Shuler et al. 1999, Averbeck and Lee 2004, Nicolelis and Lebedev 2009).

Among the studies on motor and premotor cortical neuronal populations, one paper by Churchland and his colleagues (Churchland, Cunningham et al. 2012) became especially popular. In their work, Churchland et al. claimed to have discovered a unique property of cortical population activity that they called “rotational dynamics.” Their analysis is based on the idea that motor cortical activity could be modeled as a dynamical system:

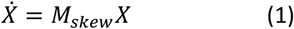

where *X* is a multidimensional vector representing neuronal population activity, 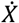 is its time derivative and *M*_*skew*_ is the transform matrix. *M*_*skew*_ has imaginary eigenvalues, which means that 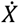 is orthogonal to *X* (Garfinkel, Shevtsov et al. 2017), which corresponds to *X* rotating with the center of rotation at the center of coordinates (Figure 1A). In the analysis of Churchland et al., vector *X* is produced from the activity of neuronal populations by the application of principal component analysis (PCA). The first six principal components (PCs) are selected to avoid overfitting that could take place with a larger number of dimensions. Hence, *X* is six-dimensional. Next, a method called jPCA is applied to compute *M*_*skew*_ and the corresponding eigenvectors. Churchland et al. projected PC data to the plane defined by the first two most prominent rotational components generated with jPCA and obtained convincing looking figures, where population responses rotated in the same direction for different experimental conditions (Figure 1B).

**Figure 1.**
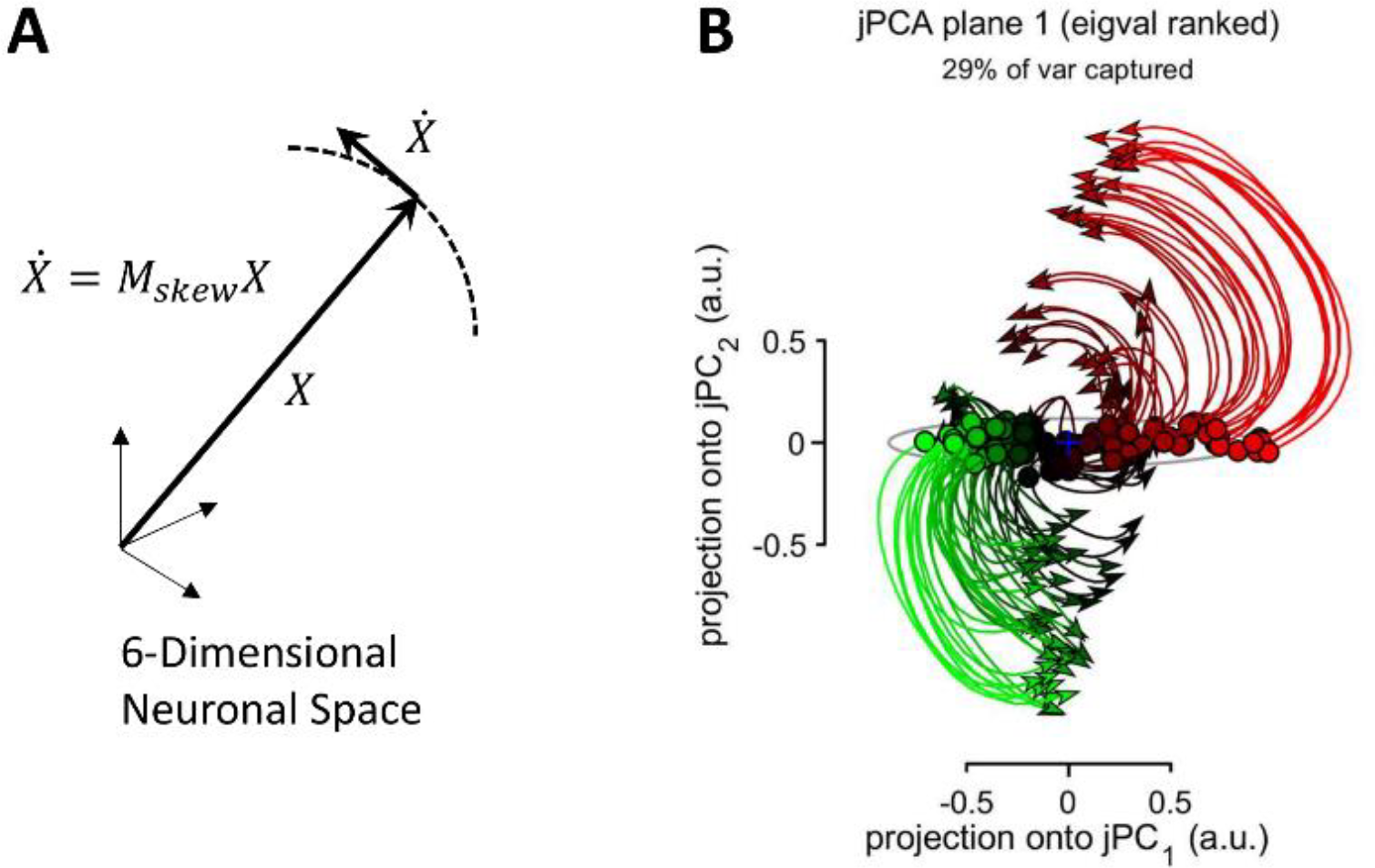
Rotation in a multidimensional neuronal space. **A:** Schematics of “rotational dynamics”, where there is an angle between the neuronal vector, *X*, and its time derivative, 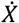, meaning that *X* is turning. In jPCA proposed by Churchland et al. (2012), is *X* is formed by the first six PC of the population activity. **B:** Application of jPCA to neuronal data. Curves represent experimental conditions, where a monkey performed armed reaching with different trajectories. The color of the curves (red, green, black) corresponds to different levels of premovement activity.

According to the interpretation of Churchland et al., “rotational dynamics” are a fundamental feature of population activity in the motor and premotor cortical areas and a proof that motor cortical populations act as a dynamical system rather than representing various motor parameters by their firing. They called the rotational effect an “orderly rotational structure, across conditions, at the population level”, a “brief but strong oscillatory component, something quite unexpected for a non-periodic behavior”, and “not trivial” observations that are “largely expected for a neural dynamical system that generates rhythmic output.”

While this proposal has merits, we found the results of Churchland et al. difficult to comprehend because of the lack of clarity regarding the concrete neuronal patterns contributing to the rotations revealed by jPCA. For example, they suggested that “motor cortex responses reflect the evolution of a neural dynamical system, starting at an initial state set by preparatory activity […]. If the rotations of the neural state […] reflect straightforward dynamics, then similar rotations should be seen for all conditions. In particular, the neural state should rotate in the same direction for all conditions, even when reaches are in opposition” – a high-level description that is hard to understand because it lacks concrete details. For instance, it is unclear what “straightforward dynamics” are and why they impose the same rotational patterns on all conditions. The major obstacle to understanding the ideas expressed by Churchland and his colleagues is the absence of a clear explanation of how individual neurons and/or their populations form the rotating patterns revealed by jPCA.

To eliminate this gap in understanding, we reanalyzed some of the data that Churchland et al. made available online. We utilized perievent time histograms (PETHs), a basic method for displaying neuronal data, to find the underlying cause for the “rotational dynamics.” Next, we ran simple simulations to verify our findings. Based on our results, we found the interpretation offered by Churchland et al. highly overstated and overreaching. Even though a certain temporal order in which different cortical neurons are activated during a behavioral task could be described as “rotational dynamics”, we are not convinced that this observation alone could significantly “help transcend the controversy over what single neurons in motor cortex code or represent,” as stated by Churchland and his colleagues.

## Results

Rotation in a multidimensional neuronal space could be thought of as a process where individual neurons are activated in a certain order, which results in the neuronal vector *X* changing orientation (Figure 1A). Such a pattern can be also described as a phase shift between the responses of different neurons. Consider the simplest case of a population that consists of two neurons where the activity of the first neuron is a sine function of time and activity of the second neuron is a cosine. Since the phase shift between the responses of these neurons is 90 degrees, a two-dimensional plot with the firing rates of these neurons on the axes produces circular or elliptical trajectories. This type of trajectory is observed for all conditions if the phase shift between the neurons persists.

Following this logic, we hypothesized that the data of Churchland et al. contained phase shifts between the neurons, which remained consistent across conditions. To test this hypothesis, we analyzed their data using the traditional PETH method. The dataset included PETHs of 218 neurons calculated for 108 experimental conditions (i.e., one smoothed PETH per condition; single-trial data were unavailable). Each condition corresponded to a monkey performing a reaching movement with a straight or convoluted trajectory. We simply stacked these PETHs to produce population color plots for different conditions (Figure 2A-D). Additionally, we averaged the PETHs across all conditions to obtain average responses for each neuron (Figure 2E). For the average PETHs, we calculated peak values and reordered the neurons according to the value of the time when each neuron reached its peak firing rate. In the color plot showing the average PETHs (Figure 2E), PETHs of the neurons activated early are plotted at the top and PETHs of the neurons activated late are plotted at the bottom, which results in a clear display of an activity wave running across the neurons in the population. The same reordering was applied to the PETHs for individual conditions (Figure 2A-D). In the PETHs for individual conditions, the temporal order of responses persisted with some jitter (e.g., compare panels A, B, C and D in Figure 2). The same sequence of responses is also clear in the scatterplot that displays the time of peak response for different conditions and different neurons (Figure 2F). Additionally, pairwise correlations were strong for the neurons with similar occurrences of response and weak (or negative) for the neurons with dissimilar occurrences (Figure 2G). Thus, the PETH analysis showed that in Churchland’s data neurons responded in a certain temporal order, and this order persisted even when the monkey altered the way it performed a reaching movement.

**Figure 2.**
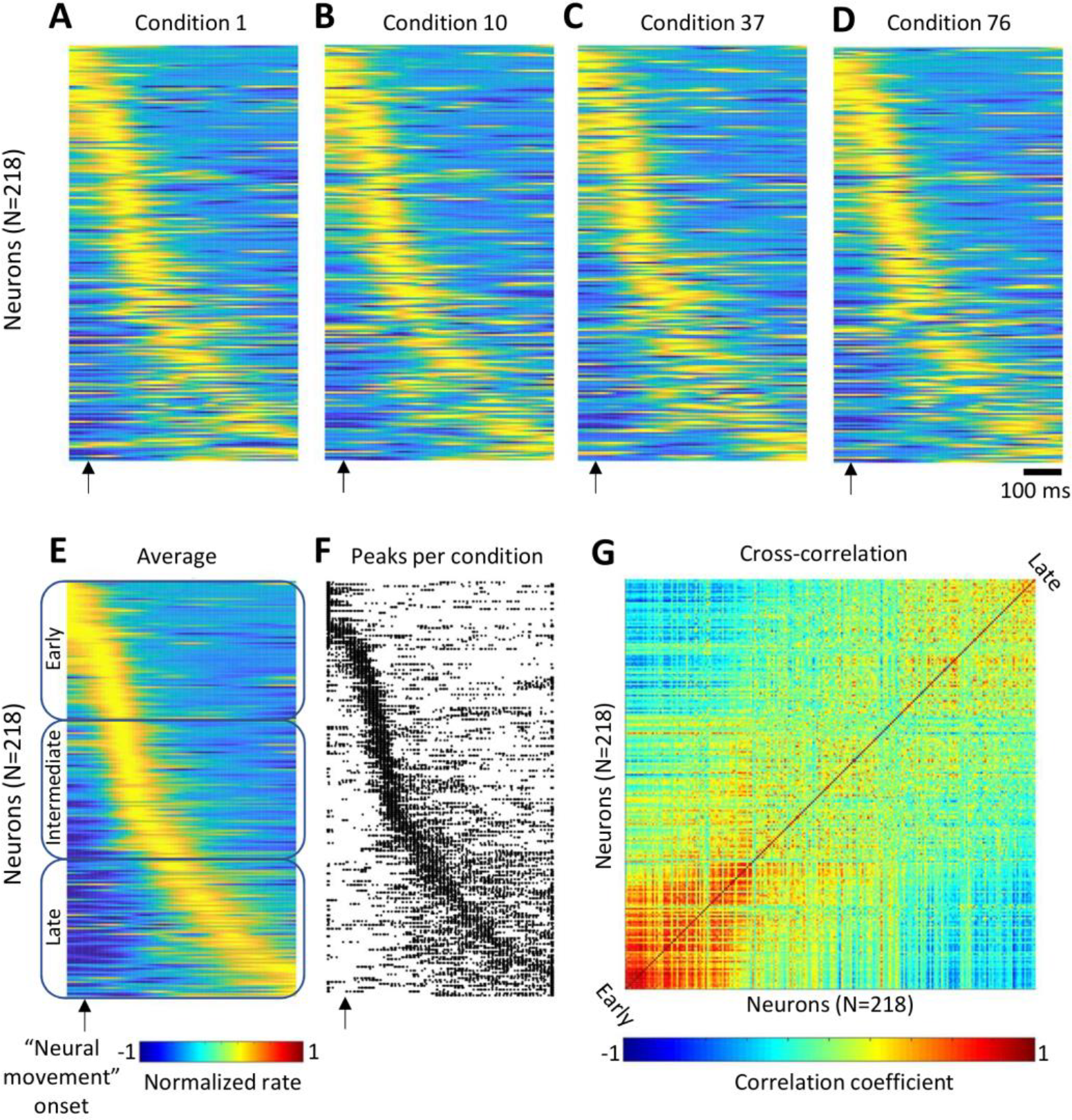
Time spread in peak firing rates of different neurons in the data provided by Churchland et al. (2012). **A-D:** Color-coded population PETHs for four representative conditions. Horizontal lines represent PETHs for individual neurons. **E:** PETHs averaged across conditions. From top to bottom: neurons are ordered by the time of peak firing rate, from the earliest activated neurons to the latest. The same order is used in A-D, F and G. The neuronal population was dived into three subpopulations: early (neurons 1-73), intermediate (74-146), and late (147-218). **F:** A scatterplot showing the times of peak firing rates for different neurons and conditions. Peak times are represented by dots. G: Pairwise correlation between the activity patterns of different neurons.

To assess how the activation order of different neurons could contribute to a population rotational pattern, we split the entire neuronal population into three subpopulations: neurons with early peaks (ranks 1-73), intermediate peaks (74-146), and late peaks (147-218) (Figure 2E). Next, we calculated average PETHs for each subpopulation and for each experimental condition (Figure 3A). As expected, this plot revealed three groups of PETHs (early-peak, intermediate-peak and late-peak) whose shapes did not change substantially across conditions. Plotting these PETHs in a three-dimensional space (where dimensions corresponded to the subpopulations) yielded a family of curved trace (Figure 3C) that resembled the circles obtained with jPCA (Figure 1B). As an additional control, we randomly shuffled conditions for each neuron and plotted the curves for the shuffled data (Figure 3B,D). Both the average PETHs for the subpopulations (Figure 3B) and the three-dimensional plot (Figure 3D) were little affected by the shuffling procedure, which confirmed that the activation order of the neurons was approximately the same for different conditions.

**Figure 3.**
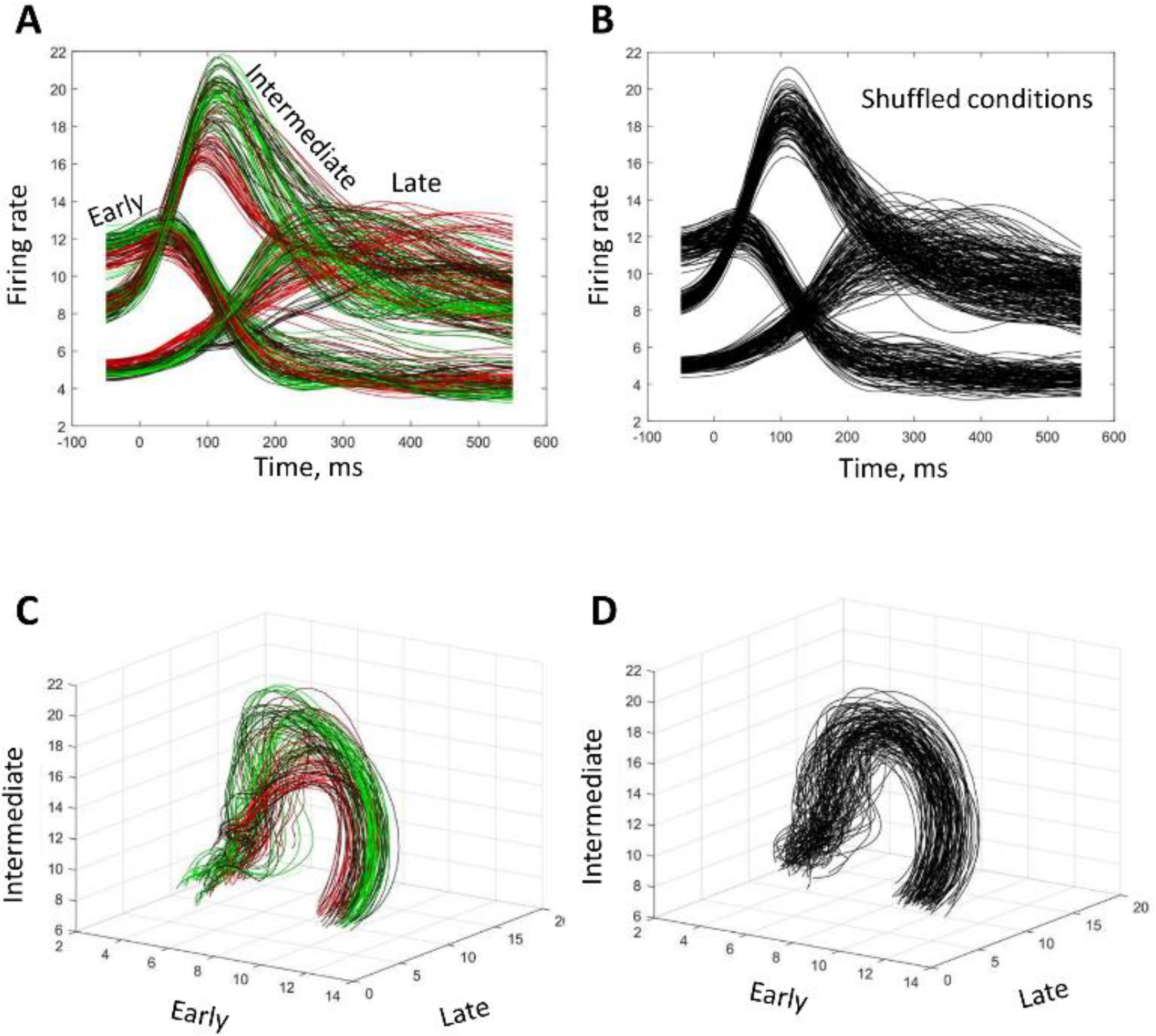
“Rotational dynamics” revealed by splitting neuronal population into the early, intermediate and late activated subpopulations. **A:** Average PETHs for the three subpopulations, for different experimental conditions. The composition of the subpopulations is shown in Figure 2E. Color coding of the traces is the same as in Figure 1B. **B:** Average PETHs for the shuffled-conditions data. **C, D:** “Rotational dynamics” shown as three-dimensional plots with the axes representing average PETHs for different subpopulations. Original (C) and shuffled (D) datasets are shown.

To further clarify the origin of the rotational patterns, we calculated the initial three PCs for the data of Churchland et al. and plotted them as a three-dimensional plot (Figure 4A). The PC traces were clearly curved. Additionally, distinct clusters of conditions were visible in the plots, each of them containing several traces that had similar shapes. The clusters started from approximately the same point but separated toward the end of the trial. Despite the differences between the clusters, they rotated in approximately the same fashion (e.g., in the plane defined by the first and second PCs). Thus, “rotational dynamics” were clearly visible even before the application of jPCA. As to JPCA, it also yielded several clusters of circles (Figure 4C).

**Figure 4.**
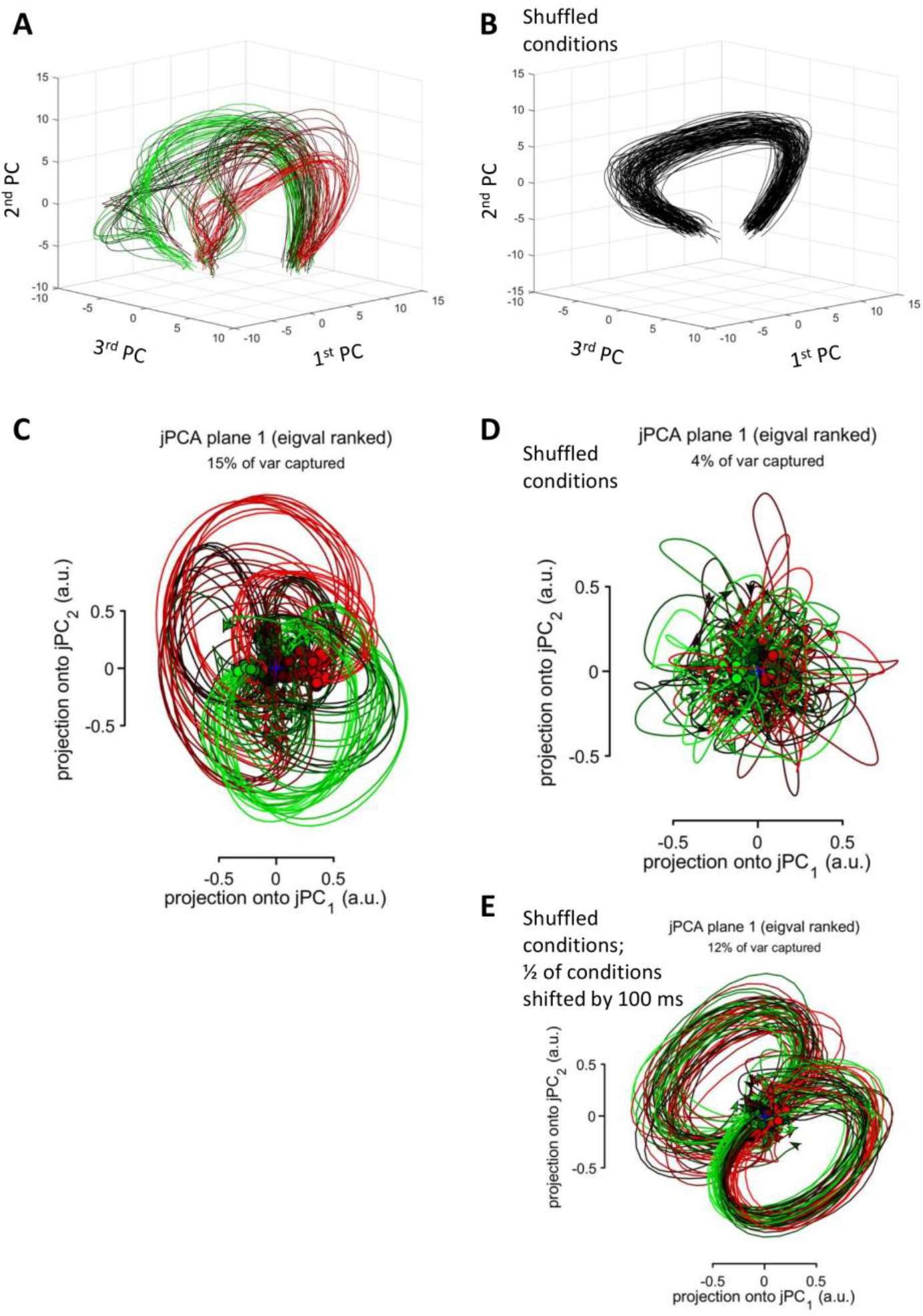
Rotations of principal components. **A:** The first three PCs plotted as a three-dimensional plot. Color conventions as in Figure 1B. **B:** PCs for the data with shuffled conditions. **C:** jPCA results for the data in A. **D:** jPCA results for the data in B. **E:** jPCA results after the shuffled data (B) was shifted forward by 100 ms for one-half of the conditions.

Rotations of PC traces remained even after experimental conditions were randomly shuffled for each neuron (Figure 4B), which indicated that condition-specific correlation between the responses of different neurons was not crucial for a rotational pattern to occur. While the PC traces for the shuffled data were clearly curved, they were not clustered any more after the shuffling procedure. Unlike PCA results, jPCA (with the default settings, but see Figure 8 for jPCA results with different settings) failed to detect these obvious curvatures and returned noisy traces (Figure 4D). This result indicated that, after the shuffling procedure made the population responses very similar for different conditions, the data became ill-conditioned (Golub and Van Loan 2012) for jPCA. We, however, found a simple solution to this problem. We introduced variability to the entries for different conditions: for half of the conditions (conditions 55-108), the PETHs of all neurons were shifted forward by 100 ms, and the initial 100 ms of the PETHs were assigned constant values that were equal to the first point of each original PETH. Such activity pattern would have occurred if the monkey delayed movement initiation by 100 ms. Importantly, this manipulation did not alter the temporal sequence of neuronal population. Following this shift, jPCA returned circles (Figure 4E).

As a side note, jPCA with default settings encounters the same problem for a simple simulation where responses of a half of the neurons are a sine function and responses of the other half are a cosine. Although this is an obvious case of “rotational dynamics,” jPCA fails to generate circles. However, the circles appear after the population responses are shifted in time by a random amount for different conditions. As explained below, the problem of ill-conditioning occurs because jPCA incorporates by default a data preprocessing step where, for each neuron, an across-condition average PETH is subtracted from all PETHs for that neuron.

Having established that neurons were activated in consistent temporal order in Churchland’s data, we examined this effect further using a simple simulation of population activity. This simulation did not incorporate any features specific for motor cortical responses. For example, simulated neurons were not directionally tuned and instead each of them exhibited random amplitude of response for different conditions. Because of this randomness, response amplitude was not correlated in any pair of neurons. The only nonrandom pattern that we simulated was the presence of a sequence of responses in different neurons. The shape of simulated PETHs was a Gaussian function with an amplitude drawn from a uniform distribution (Figure 5). We simulated 218 neurons and 108 conditions to match Churchland’s data.

**Figure 5.**
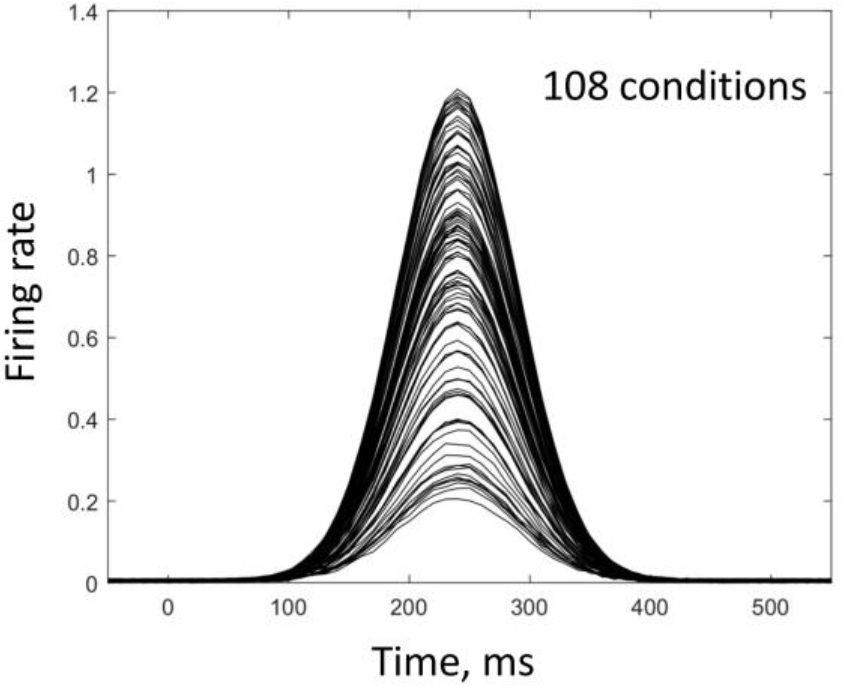
Simulated responses for an individual neuron. Neuronal responses were simulated using a Gaussian function. Response amplitude was randomly drawn from a uniform distribution (in the interval 0.2-1.2).

Recently, Michaels, Dann et al. (2016) used a somewhat similar simulation of motor cortical neurons with a temporal sequence of responses that was consistent across conditions. They observed an occurrence of “rotational dynamics” under these conditions. Their simulation was, however, more sophisticated than ours as they simulated directionally tuned motor cortical neurons and attempted to compare a representational model of population activity with a dynamical-system model. To prove that jPCA reveals “complex aspects” of motor cortical activity, they designed sophisticated methods for data permutations. Yet, they did not consider the possibility that some of these permutations made their simulated data ill conditioned for jPCA to return circular trajectoris. In our simulation, resemblance to motor cortical activity was minimal. We also applied a simple correction to cope with the ill conditioned data. Moreover, our overall conclusion is the opposite to the opinion Dann et al. (2016) who highly valued jPCA approach and used it as the gold standard for assessing the results of their simulation.

We started with a simulation, where the responses of different neurons were shifted in time: neuron *i* responded 1 ms later than neuron *i-1* (Figure 6A). This simulation produced a wave of activity running through the neuronal population. Here, the population responses were very similar for different conditions as in the results for shuffled data presented above. Apparently, in this case the data was ill-conditioned for jPCA, so this analysis returned noise (Figure 6B) even though the input data contained a consistent temporal sequence of neuronal responses. To make neuronal responses more variable across conditions and thereby remove the ill conditioning, we simulated two groups of conditions: for conditions 1-54, the first neuron exhibited peak activity at time *t* = 50ms, and for conditions 55-108 the time of its peak activity was *t* = 200ms (Figure 6C). The structure of the activity wave (i.e. the rule that neuron *i* responded 1 ms later than neuron *i-1*) was the same for both groups of trials. This slight alteration of the data, which did not change the structure of the population response, was sufficient for jPCA to start generating circles (Figure 6D). We also simulated three groups of conditions, where the first neuron’s peak rate occurred at 50, 150 and 200 ms for the first, second and third groups, respectively (Figure 6E). The structure of the population wave was the same for all three groups. In this case, again, jPCA returned circles (Figure 6F).

**Figure 6.**
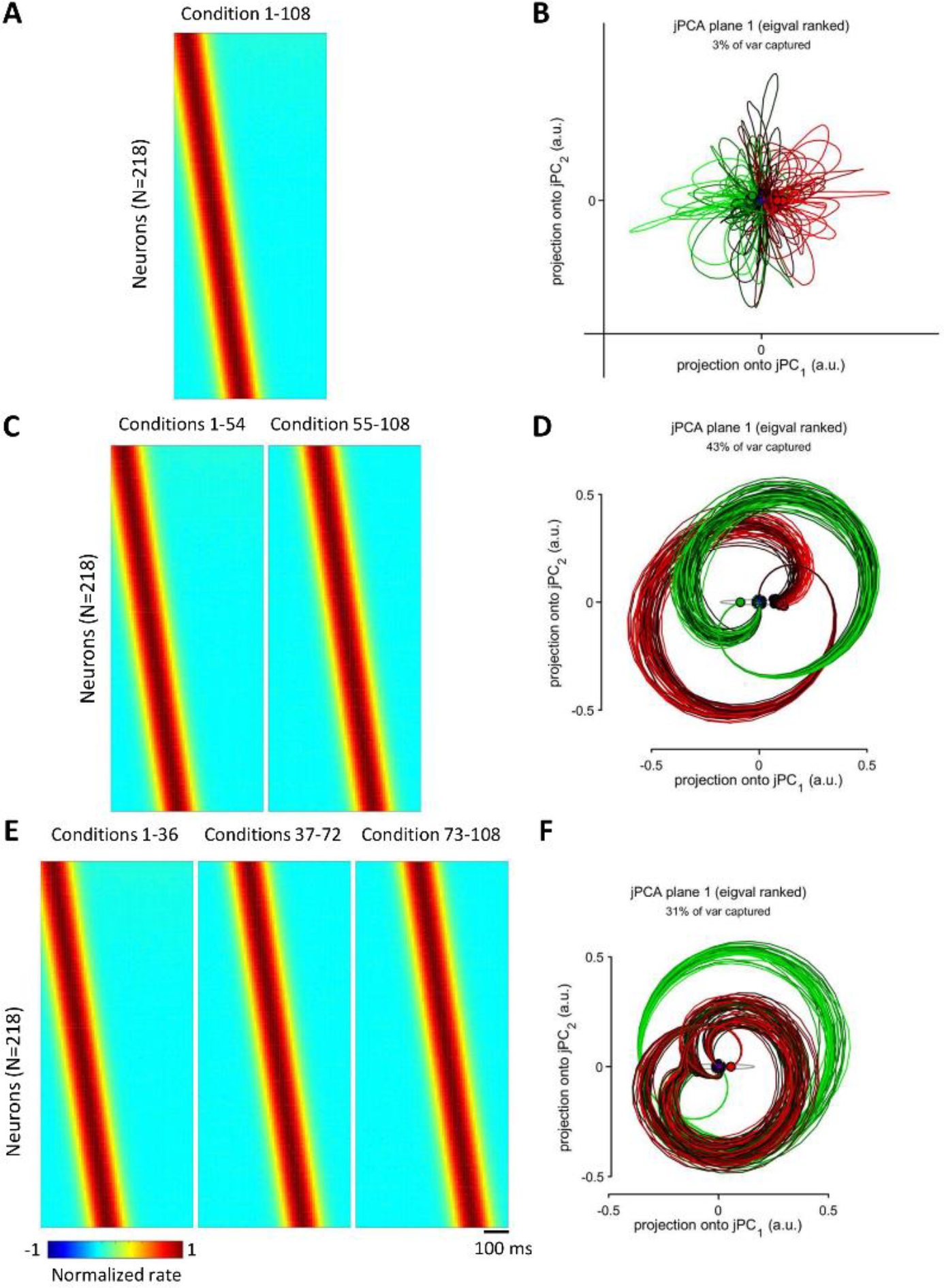
Simulated population responses with a time spread of peak firing rates. **A:** PETHs for a simulated wave of population activity, where response of neuron *i* occurs 1 ms later than response of neuron *i-1*. **B:** jPCA results for the data in A. **C:** The same population wave as in A with an early onset for conditions1-54 and late onset for conditions 55-108. **D:** jPCA results for the data in C. **E:** The same population wave as in A and C with three different onsets for conditions 1-36, 37-72 and 73-108. **F:** jPCA results for the data in E.

To verify that a temporal sequence of activation was necessary for the rotation pattern to occur for our simple model of neuronal responses, we simulated a neuronal population where all neurons responded simultaneously (Figure 7). The shapes of neuronal responses were the same as in the previous simulation (Figure 5). In this case, jPCA failed to generate circles (Figure 7B) even when three groups of conditions were simulated with shifted activity onsets (Figure 7A).

**Figure 7.**
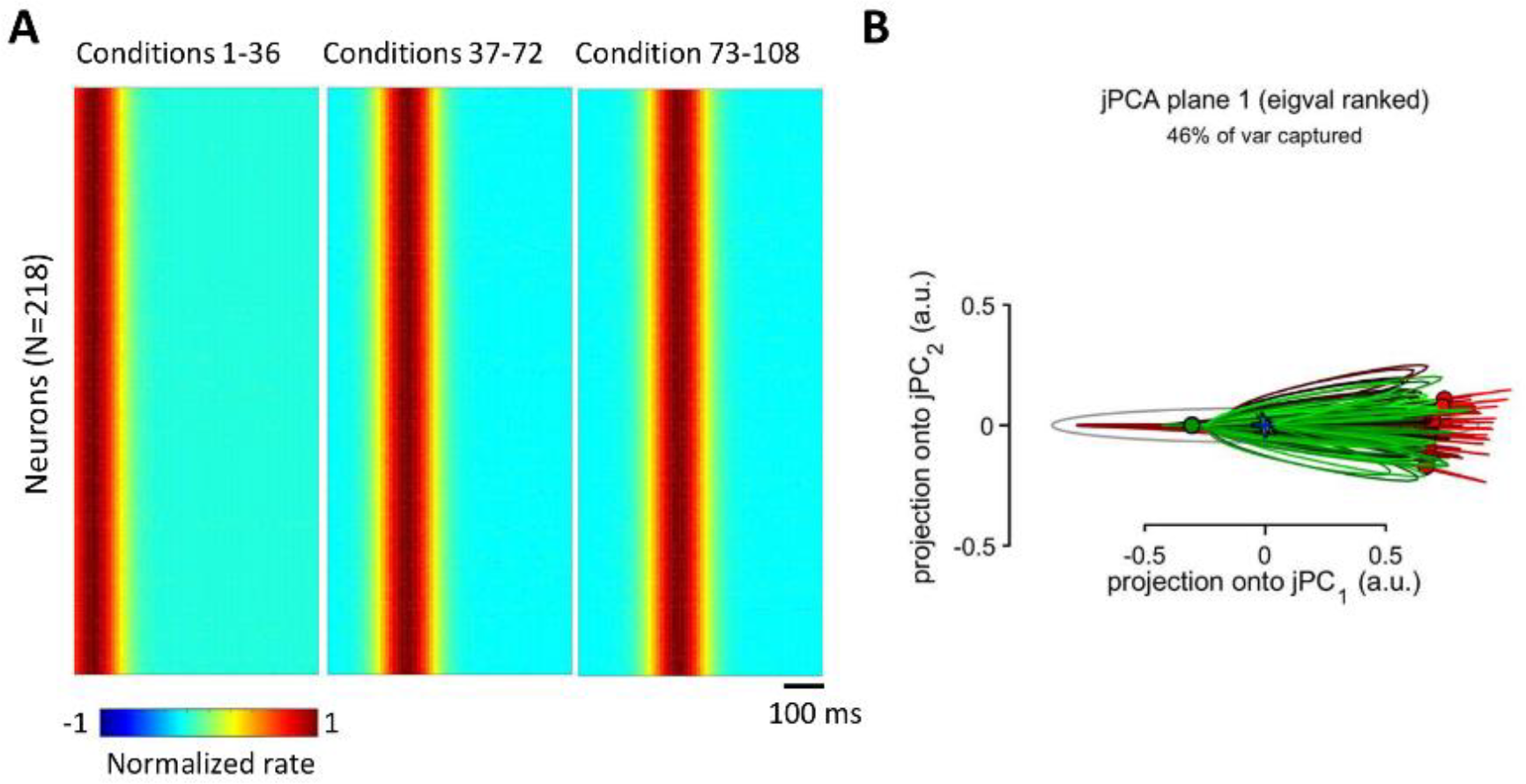
Simulated population responses without a time spread of peak firing rates. **A:** PETHs with different response onsets for conditions 1-36, 37-72 and 73-108. For each group of conditions, peak responses of different neurons were not spread in time. **B:** jPCA results for the data in A.

In addition to running jPCA with the default settings, where an average across-condition response was subtracted from each PETH, we ran jPCA without this setting (Figure 8). In this case, the problem ill conditioning did not occur, and jPCA returned circles in all cases with the exception of the simulation, where there was no temporal sequence of neuronal responses (Figure 8D). Overall, probing our simulated data with different versions of Churchland’s jPCA supported the hypothesis that a temporal sequence of neuronal responses results in a population rotation pattern.

**Figure 8.**
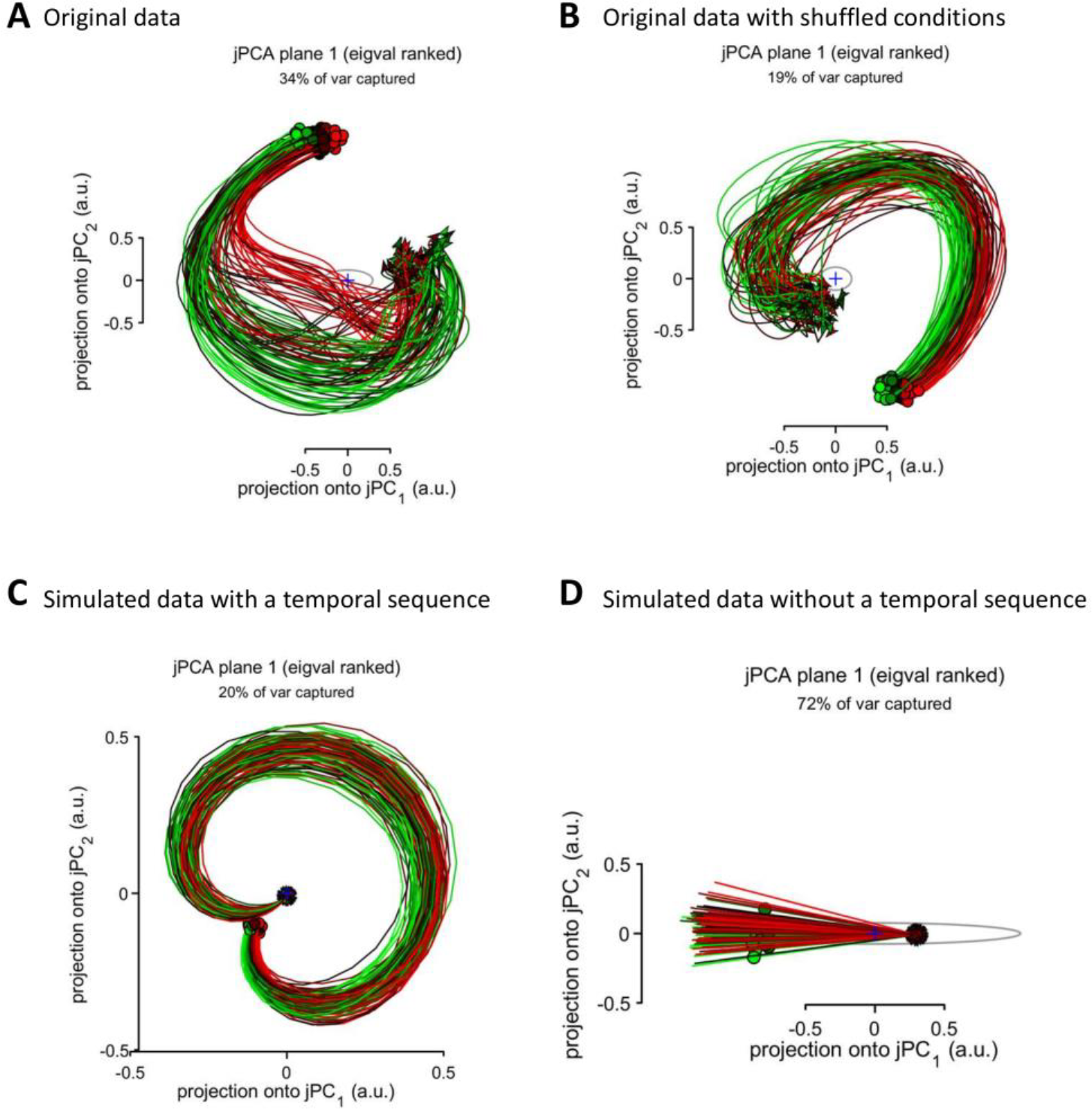
jPCA results without subtracting across-condition average responses. **A:** Original data of Churchland et al., the same data as in Figure 4C. **B:** Original data after across-condition shuffling, the same data as in Figure 4D. **C:** Simulated data with a temporal sequence of responses, the same data as in Figure 5A,B. **D:** Simulated data without a temporal sequence of responses, the same data as in Figure 7.

Finally, we probed a more direct approach for converting Churchland’s data into circular trajectories. Churchland et al. linearly fit the population activity vector to its first derivative (equation 1) with the goal of extracting a rotational structure. We hypothesized that such an extraction of a desired “population property” could be achieved with a multiple linear regression. Accordingly, we utilized multiple linear regression to fit neuronal data to a circle:

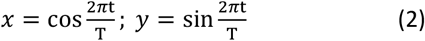

or to the Lissajous curve shaped as ∞:

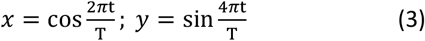

where *t* is time and *T* is trial duration. The regression worked well for both shapes (Figure 9A,B), and it even generalized to new conditions, as evident from the analysis where half of the data were used for training the regression model and the other half for generating predictions (Figure 9C,D). The shuffled data could be fit to the Lissajous curves, as well (Figure 9E-H), although the prediction of ∞ was very noisy (Figure 9H).

## Discussion

To clarify the neurophysiological significance of “rotational dynamics,” we conducted additional analyses of Churchland’s data and performed simple simulations. In the original dataset, we discovered a temporal order in which the neurons were activated and found that this order was relatively consistent across conditions. In our opinion, this is a useful observation that has not been described in sufficient detail in the original article of Churchland et al. and the publications that emerged from this study. We suggest that simple plots of population PETHs (e.g., Figure 2) should be used to clarify any results on “rotational dynamics”. These plots could be very useful to clarify claims like the “rotational dynamics” persist even though motor-related patterns are different across condition (Churchland, Cunningham et al. 2012) or the recurrent neural network trained to generate arm-muscle EMGs develops rotational patterns (Sussillo, Churchland et al. 2015).

**Figure 9.**
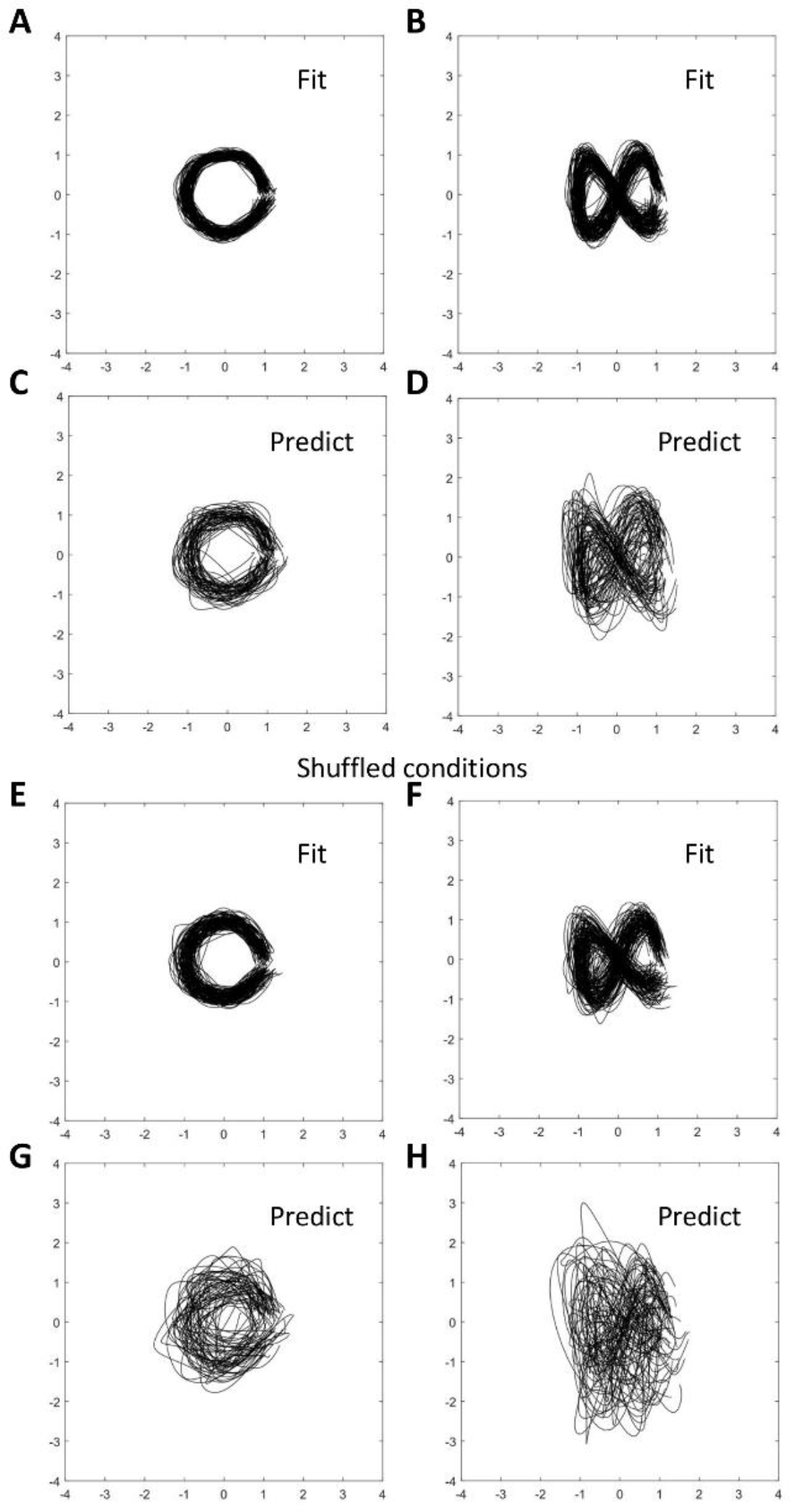
Fitting population data to Lissajous curves. **A:** Fitting to a circle. **B:** Fitting to the curve shaped as ∞. **C:** Prediction of a circle. Half of the conditions were used to train the regression model; the other half to generate prediction. **D:** Prediction of the ∞ shape. **E-H:** Fitting and predicting for the data with shuffled conditions.

The existence of a temporal spread in neuronal peak rates (or response latencies) has been extensively documented in the literature. For example, waves of activity similar to those shown in Figure 2E-F have been depicted in numerous publications (Luczak, Barthó et al. 2007, Gage, Stoetzner et al. 2010, Peyrache, Benchenane et al. 2010, Kvitsiani, Ranade et al. 2013, Bulkin, Law et al. 2016). As to motor cortical patterns during reaching, Figure 7 in Georgopoulos et al. (Georgopoulos, Kalaska et al. 1982) shows a distribution of the times of onset of the first increase in discharge in motor cortical neurons for center-out arm reaching movements performed by a monkey. The onsets were calculated for the neurons’ preferred directions and ranged −300 to 500 ms relative to movement initiation time. In a more recent paper (Ifft, Lebedev et al. 2011), we have demonstrated such a spread for simultaneously recorded populations of neurons recorded in the motor and primary somatosensory cortical areas.

It may be true that no previous paper focused on the consistency of neuronal activation order for a range of movements. Yet, Churchland and his colleagues did not emphasize such consistency either and instead emphasized variability of neuronal responses across conditions. According to their explanation, rotational patterns are principally a population phenomenon that persists despite individual neurons exhibiting different activity patterns for different arm reaches.

Temporal sequences of neuronal responses that are consistent across a set of experimental conditions could occur for various reasons. For example, such a temporal order could be related to serial information processing by a neuronal circuit. Orderly response onsets have been reported for the activity of multiple cortical areas transforming sensory inputs into behavioral actions. For example, de Lafuente and Romo (2006) reported that a wave of neuronal activity travelled from the somatosensory cortex to the premotor cortical areas when monkeys performed a task that required perceptual judgment of a tactile stimulus. Sensorimotor transformation of this kind could be analyzed using a comparison of response characteristics for different cortical locations and different types of neurons. Unfortunately, jPCA makes such an analysis impossible because it lumps the activity of many neurons together (e.g. compare Figure 2 with Figure 4C).

Based on the results of our analysis, we question that Churchland’s results provide a strong evidence in favor of the dynamical-system that acts “like a spring box” that “could be released to act as an engine of movement” (a comparison used by Michaels, Dann et al. (2016)). This is simply because the mere presence of a temporal sequence of neuronal responses tells us very little about the properties of the system even if we assume that it is dynamical. Temporal lags between the responses of different neurons could occur for various reasons, including direct connectivity, common input, or association with different stages of information processing. jPCA would reveal rotations even in the combined data from different monkeys – an obviously artifactual cause for lags between the responses of different neurons, and there would not be a way to tell that this is not a “dynamical system”. (Unless correlations between neurons are analyzed across behavioral trials; but such an analysis is not incorporated in the Churchland et al. approach.)

The finding that neurons in some parts of the nervous system respond to a stimulus with different delays (and jPCA reveals rotations) is not surprising as it has been published in myriads of papers. For example, in sensory systems, peripheral receptors are activated first, followed by neuronal activity in the spinal cord, brainstem, thalamus and eventually cortex. Although this sensory processing chain in principle could be defined as a dynamical system, this definition alone does not illuminate any mechanism of sensory processing. By the same token, motor systems of the brain are well-known to perform sensorimotor transformations (Georgopoulos, Lurito et al. 1989, Kalaska 1991). In Churchland’s case, this is the transformation of a go-cue and preceding instructions into a motor execution command. Saying that “rotational dynamics” occur during this sensorimotor transformation only adds terminology but does not clarify any concrete neuronal mechanism.

According to Churchland et al., “a focus on the dynamics that generate movement will help transcend the controversy over what single neurons in motor cortex code or represent.” While we agree with this general idea, we do not see how their jPCA results transcend any controversy, particularly given the fact that population rotations could be obtained with little effort using simple and quite arbitrary simulations that did not mimic any important property of motor cortical activity. Without a linkage to a specific function of the motor cortex, jPCA appears to be just a way to produce an illustration of the lags between the responses of different neurons.

Although analysis of response lags in different neurons could be valuable for understanding neural processing pipelines, we are not convinced that jPCA is the best tool for such an analysis. We observed cases where temporal sequences of responses were revealed with several methods but not with jPCA set to default, where for each neuron an across-condition response was subtracted from this neuron’s PETHs. Thus, shuffling Churchland’s data across conditions did not destroy the rotations revealed with non-jPCA approaches (Figure 3B,D and Figure 4B) but jPCA failed to produce circular trajectories in this case (Figure 4D). This happened because jPCA was designed to suppress population rotations that occurred in all conditions. This was achieved by calculating an across-condition average PETH for each neuron and then subtracting this average response from all PETH for that neuron. If this data preprocessing step was removed from jPCA, jPCA generated circles for the shuffled data (Figure 8B) and simulated temporal sequences of neuronal responses (Figure 8C). Looking closer at the preprocessing procedure employed by Churchland et al., we find it arbitrary and questionable. The first concern is that this data manipulation violates the assumptions of equation 1, where actual neuronal rates are used instead of neuronal rates contaminated by average contributions from different conditions. If Churchland et al. wanted to hypothesize that an average response influenced population dynamics, they should have incorporated this function of time in equation 1 and explained what brain structure could generate it. Unless such an explanation is provided, it is unclear why one would need to subtract an across-condition average from the neuronal response. Consider an example of a mass on a string, a harmonic oscillator that could be described as a dynamical system (Hirsch, Smale et al. 2012). It would make little sense to analyze the properties of this system by subtracting average responses for different conditions like different values of mass and spring constant, which influence the period of oscillations. (This example is relevant to motor control where biomechanical parameters are different for different movements.) Calculation of an across-condition average is an artificial procedure because it requires aligning neuronal data on an event of interest, for example go-cue onset, movement onset, or Churchlan’s neuronal movement onset. For data points distant from this aligning event, variability in neuronal responses would grow (Renoult, Roux et al. 2006), making the average PETH a less reliable description of neuronal activity as a function of time and, consequently, an ineffective factor for Churchland’s data correction. Additionally, this correction cannot be applied to continuous behaviors like target pursuit task (Li, O’Doherty et al. 2009), where phase differences between the activity of different neurons are very likely to occur but the task is not composed of epochs corresponding to a discrete set of conditions. Churchland and his colleagues justified their correction procedure by the desire to remove “rotations that are similar for all conditions” they prevent one “to gain multiple views of the underlying process, making it difficult to infer whether rotations are due to dynamics or to more trivial possibilities”. While they did not clarify the difference between the “dynamics” and “more trivial possibilities”, the consequences of the implemented data manipulation are not trivial. It may have caused aliasing, such as significant distortion of neuronal responses. Overall, this manipulation appears poorly justified and its benefits are unclear.

Mathematically, Churchland’s method to produce a rotating pattern from neuronal PETHs can be described as a linear transformation with a certain number of free parameters. With a sufficient number of free parameters, multiple linear regression can yield practically any desired curve (Babyak 2004). For example, when we applied a multiple linear regression to Churchland’s data, we easily produced a variety of Lissajous curves. Because of the similarity of neuronal responses across conditions, this transformation even generalized from the training dataset (half of the conditions) to new data in the test dataset (the other half) (Figure 8). Evidently, no one would claim that any of these arbitrarily chosen curves represents a physiologically meaningful neural population mechanism, although we used a linear transformation very similar to the one Churchland and his colleagues employed to generate their curves.

Altogether, our results challenge the conclusions of Churchland and his colleagues. Clearly, their effects simply reflect what one should expect by applying a linear transformation, such as PCA, to reduce the dimensionality of large-dimension data and then adjusting the resulting components to a new basis that reproduces a desired behavior (in this case rotation). PCA (Chapin and Nicolelis 1999) and independent component analysis (Laubach, Shuler et al. 1999) were introduced twenty years ago by our group to extract correlated neuronal patterns and reduce dimensionality of neuronal-ensemble data. While the findings of Churchland and his colleagues can be viewed as an extension of this approach and a method to produce a visually appealing phase plot, we doubt that they have discovered any new neurophysiological mechanism, as claimed in their article. Indeed, the mere fact that neurons in a population respond at different times with respect to a go-cue tells very little about the function of these responses and does not discard any of the previously proposed representational interpretations, such as the suggestion that neuronal populations perform a sensorimotor transformation (Georgopoulos, Lurito et al. 1989, Kalaska 1991, Kakei, Hoffman et al. 2003) or handle specific motor parameters (Georgopoulos, Kalaska et al. 1982, Georgopoulos, Ashe et al. 1992, Kakei, Hoffman et al. 1999, Zhuang, Lebedev et al. 2014).

Cortical activity is dynamical in the sense that neuronal rates change in time. Differential equations (Lukashin and Georgopoulos 1993) and other mathematical methods (Kawato and Wolpert 1998, Moody and Zipser 1998, Todorov 2000) are certainly applicable to modeling these dynamics. However, a much more thorough analysis compared to the method proposed by Churchland et al. is needed for making a major advance in our understanding of cortical mechanisms of motor control.

## Methods

The data (single units and good multiunits recorded in monkey N; the data used to construct Fig. 3f of Churchland et a.) and MATLAB scripts were obtained from Churchland lab’s online depository (https://www.dropbox.com/sh/2q3m5fqfscwf95j/AAC3WV90hHdBgz0Np4RAKJpYa?dl=0). The dataset contained PETHs for each neuron and each condition. We used the following commands to run Churchland’s code:

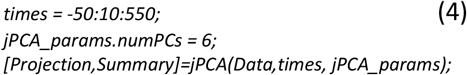

This corresponds to the time range −50 to 550ms and six PCs entered in jPCA.

To produce the plots shown in Figure 2A-D, we stacked Churchland’s PETHs together, and Figure 2E shows PETHs averaged across conditions. The average PETHs were used to find peak firing rates and the time of their occurrences. PETHs of Figure 2A-E were sorted according to the sequence of these peaks of the average PETHs. To improve the display of phase shifts between the neurons, PETHs of Figure 2A-E were normalized by subtracting the mean and dividing by the peak PETH value. This normalization was used only for plotting the graphs of Figure 2A-E but not for calculating average PETHs for individual neurons (Figure 2E) or neuronal subpopulations (Figure 3).

In the PCA analysis (Figure 4), we standardized PETHs for each neuron by subtracting the overall mean (i.e., average for all conditions combined) and dividing by the overall standard deviation (again, for all conditions combined).

Simulated PETHs (Figure 5) were computed in MATLAB as:

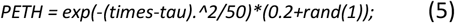

where the time shift, *tau*, was selected to produce 10-ms increments of the delay for the neurons in the sequence (Figure 6). Neither the width of the response not the amplitude (uniformly distributed from 0.2 to 1.2 in equation 5) were critical for the rotations to occur. However, to cope with the ill-conditioning of the population responses highly correlated across conditions, it was important to introduce temporal variability to the simulated PETHs. This was done by offsetting *tau* for all neurons by the same amount of time for several groups of conditions (Figure 6C,D).

Multiple linear regressions (Figure 9) were implemented in MATLAB (*regress* function). Here, neuronal activity was transformed into Lissajous curves. Fitting (Figure 9A,B,E,F) was conducted by using the same conditions as the training and test data. Predictions (Figure 9C,D,G,H) were computed by using half of the trials to train the regression model and the other half to test.

## References

Averbeck, B. B. and D. Lee (2004). “Coding and transmission of information by neural ensembles.” Trends in neurosciences 27(4): 225–230.

Babyak, M. A. (2004). “What you see may not be what you get: a brief, nontechnical introduction to overfitting in regression-type models.” Psychosom Med 66(3): 411–421.

Bulkin, D. A., L. M. Law and D. M. Smith (2016). “Placing memories in context: Hippocampal representations promote retrieval of appropriate memories.” Hippocampus 26(7): 958–971.

Chapin, J. K. and M. A. Nicolelis (1999). “Principal component analysis of neuronal ensemble activity reveals multidimensional somatosensory representations.” Journal of neuroscience methods 94(1): 121–140.

Churchland, M. M., J. P. Cunningham, M. T. Kaufman, J. D. Foster, P. Nuyujukian, S. I. Ryu and K. V. Shenoy (2012). “Neural population dynamics during reaching.” Nature 487(7405): 51–56.

de Lafuente, V. and R. Romo (2006). “Neural correlate of subjective sensory experience gradually builds up across cortical areas.” Proceedings of the National Academy of Sciences 103(39): 14266–14271.

Evarts, E. V. (1972). “Activity of motor cortex neurons in association with learned movement.” Int J Neurosci 3(3): 113–124.

Gage, G. J., C. R. Stoetzner, A. B. Wiltschko and J. D. Berke (2010). “Selective activation of striatal fast-spiking interneurons during choice execution.” Neuron 67(3): 466–479.

Garfinkel, A., J. Shevtsov and Y. Guo (2017). Modeling life: the mathematics of biological systems, Springer.

Georgopoulos, A. P., J. Ashe, N. Smyrnis and M. Taira (1992). “The motor cortex and the coding of force.” Science 256(5064): 1692–1695.

Georgopoulos, A. P., J. F. Kalaska, R. Caminiti and J. T. Massey (1982). “On the relations between the direction of two-dimensional arm movements and cell discharge in primate motor cortex.” J Neurosci 2(11): 1527–1537.

Georgopoulos, A. P., J. T. Lurito, M. Petrides, A. B. Schwartz and J. T. Massey (1989). “Mental rotation of the neuronal population vector.” Science 243(4888): 234–236.

Golub, G. H. and C. F. Van Loan (2012). Matrix computations, JHU press.

Hirsch, M. W., S. Smale and R. L. Devaney (2012). Differential equations, dynamical systems, and an introduction to chaos, Academic press.

Ifft, P., M. Lebedev and M. A. Nicolelis (2011). “Cortical correlates of Fitts’ law.” Frontiers in integrative neuroscience 5: 85.

Kakei, S., D. S. Hoffman and P. L. Strick (1999). “Muscle and movement representations in the primary motor cortex.” Science 285(5436): 2136–2139.

Kakei, S., D. S. Hoffman and P. L. Strick (2003). “Sensorimotor transformations in cortical motor areas.” Neuroscience research 46(1): 1–10.

Kalaska, J. (1991). Reaching movements to visual targets: Neuronal representations of sensori-motor transformations. Seminars in Neuroscience, Elsevier.

Kawato, M. and D. Wolpert (1998). “Internal models for motor control.” Sensory guidance of movement 218: 291–307.

Kvitsiani, D., S. Ranade, B. Hangya, H. Taniguchi, J. Huang and A. Kepecs (2013). “Distinct behavioural and network correlates of two interneuron types in prefrontal cortex.” Nature 498(7454): 363.

Laubach, M., M. Shuler and M. A. Nicolelis (1999). “Independent component analyses for quantifying neuronal ensemble interactions.” Journal of neuroscience methods 94(1): 141–154.

Li, Z., J. E. O’Doherty, T. L. Hanson, M. A. Lebedev, C. S. Henriquez and M. A. Nicolelis (2009). “Unscented Kalman filter for brain-machine interfaces.” PloS one 4(7): e6243.

Luczak, A., P. Barthó, S. L. Marguet, G. Buzsáki and K. D. Harris (2007). “Sequential structure of neocortical spontaneous activity in vivo.” Proceedings of the National Academy of Sciences 104(1): 347–352.

Lukashin, A. V. and A. P. Georgopoulos (1993). “A dynamical neural network model for motor cortical activity during movement: population coding of movement trajectories.” Biological cybernetics 69(5-6): 517–524.

Michaels, J. A., B. Dann and H. Scherberger (2016). “Neural population dynamics during reaching are better explained by a dynamical system than representational tuning.” PLoS computational biology 12(11): e1005175.

Moody, S. L. and D. Zipser (1998). “A model of reaching dynamics in primary motor cortex.” Journal of Cognitive Neuroscience 10(1): 35–45.

Nicolelis, M. A., D. Dimitrov, J. M. Carmena, R. Crist, G. Lehew, J. D. Kralik and S. P. Wise (2003). “Chronic, multisite, multielectrode recordings in macaque monkeys.” Proceedings of the National Academy of Sciences 100(19): 11041–11046.

Nicolelis, M. A. and M. A. Lebedev (2009). “Principles of neural ensemble physiology underlying the operation of brain–machine interfaces.” Nature reviews neuroscience 10(7): 530.

Peyrache, A., K. Benchenane, M. Khamassi, S. Wiener and F. Battaglia (2010). “Sequential reinstatement of neocortical activity during slow oscillations depends on cells’ global activity.” Frontiers in Systems Neuroscience 3(18).

Renoult, L., S. Roux and A. Riehle (2006). “Time is a rubberband: neuronal activity in monkey motor cortex in relation to time estimation.” European journal of Neuroscience 23(11): 3098–3108.

Schwarz, D. A., M. A. Lebedev, T. L. Hanson, D. F. Dimitrov, G. Lehew, J. Meloy, S. Rajangam, V. Subramanian, P. J. Ifft and Z. Li (2014). “Chronic, wireless recordings of large-scale brain activity in freely moving rhesus monkeys.” Nature methods 11(6): 670.

Sussillo, D., M. M. Churchland, M. T. Kaufman and K. V. Shenoy (2015). “A neural network that finds a naturalistic solution for the production of muscle activity.” Nature neuroscience 18(7): 1025.

Todorov, E. (2000). “Direct cortical control of muscle activation in voluntary arm movements: a model.” Nature neuroscience 3(4): 391.

Zhuang, K. Z., M. A. Lebedev and M. A. Nicolelis (2014). “Joint cross-correlation analysis reveals complex, time-dependent functional relationship between cortical neurons and arm electromyograms.” J Neurophysiol 112(11): 2865–2887.

